# A universal spectrum annotator for complex peptidoforms in mass spectrometry-based proteomics

**DOI:** 10.1101/2025.01.18.633732

**Authors:** Douwe Schulte, Rien Leuvenink, Shelley Jager, Albert J. R. Heck, Joost Snijder

## Abstract

Accurate and comprehensive peptide spectrum annotation is a crucial step to interpret mass spectrometry-based proteomics data. While peak assignment in peptide fragmentation spectra is central to a broad range of proteomics applications, current tools tend to be specialized to a specific task. Here we present a more comprehensive tool, the Annotator, which unifies spectrum annotation for bottom-up, middle-down, top-down, cross-linked, and glycopeptide fragmentation mass spectra, from all fragmentation methods, including all ion types: a/b/c, x/y/z, d/v/w, and immonium ions. The Annotator integrates all known post-translational modifications from common databases and additionally allows for the definition of custom fragmentation models and modifications. Modifications allow for diagnostic fragment ions, site specific neutral losses, and multiple breakage sites for cross-linkers. The underlying library used for the theoretical fragmentation and matching is based on the unified peptidoform notation ProForma 2.0 and made available as a Rust library (rustyms) with Python bindings. This enables spectrum annotation of diverse and complex peptidoforms across the broad range of mass spectrometry-based proteomics applications.

## Introduction

Mass spectrometry-based proteomics encompasses a broad range of methods to study proteins^1–3^. Its applications span from bottom-up proteomics on shorter peptides to top-down proteomics of intact proteins with lengths of up to hundreds of amino acids incorporating complex disulfide linkages. Additionally, mass spectrometry is a pivotal method to study protein post translational modifications (PTMs)^4^, like phosphorylation, glycosylation and ubiquitination. The accuracy of peptide and protein identification, *de novo* sequencing, and mapping of PTMs crucially depends on the available evidence in fragmentation spectra. While many tools exist to annotate and score such spectra for the presence of the most common peptide backbone fragments^5–18^, much can be gained by extending the experiments and interpretation to a broader range of activation methods and fragmentation pathways, including diagnostic ions for specific peptide structures^19^.

The primary products of peptide fragmentation correspond to breaks at the three unique positions along the backbone, generating the well-known series of a/x-, b/y-, and c/z-ions, in which a, b, c and x, y, z represent the complementary N- and C-terminal counterparts of each position along the peptide^20^ (see Figure 1A). These fragments are characteristic of the sequence of amino acids. The occurrence and abundance of specific fragment ions depends on the activation method used (e.g. CID, ETD, UVPD), but also on the charge, sequence, and resulting chemical properties of the peptide. Backbone fragments are quite often accompanied by characteristic neutral losses such as water (−H_2_O), ammonia (−NH_3_), or in the case of electron-based fragmentation also -CHO_2_ and –C_2_H_3_O_2_ ^21,22^. Additionally, secondary fragmentation can occur, resulting in the (partial) loss of side chains to generate d-, v-, and w-ions ^23^. While the v-ion results from loss of a complete side-chain^24^, d- and wions result from cleavage at the sidechain’s C_β_ atom to generate informative fragments to distinguish, for instance, isoleucine from leucine residues^25–27^. Secondary fragments can also occur at additional backbone positions, resulting in internal peptide fragments and an explosion of theoretically possible matches in the spectrum (which has attracted special attention in top-down proteomics as of late)^28–31^. A special case of the internal fragment is the generation of immonium ions, following subsequent a- and y-type fragmentation at the same residue, providing fragments in the lower mass range (<200 Da) that inform about the amino acid composition of the peptide^32^.

**Figure 1.**
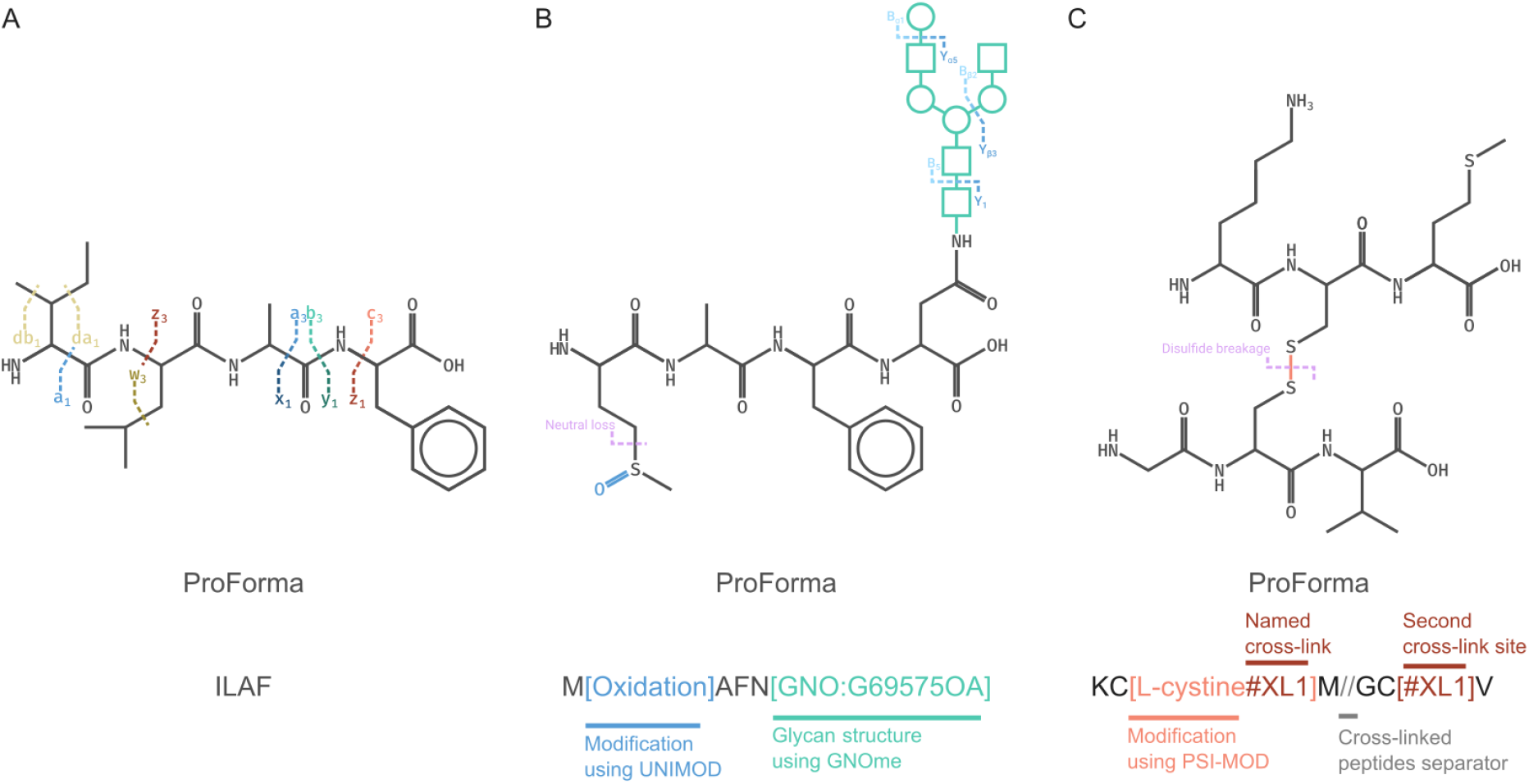
Peptidoform expression and fragment annotation in rustyms and Annotator. A) Peptides are expressed in ProForma to define their possible backbone fragments and satellite ions. B) Post-translational modifications including glycosylation are accommodated following public databases or custom definitions with diagnostic ions and neutral losses. C) Cross-linked pairs of peptides can be defined using any crosslinker, with diagnostic ions for cleavable crosslinkers, including disulfide bonds.

PTMs are often accompanied by distinct diagnostic ions, which may provide crucial evidence for the existence and localization of the modification (see Figure 1B). Common examples are that oxidation on methionine has a specific neutral loss of -C_1_H_4_O_1_S_1_ and a phosphorylated serine, threonine, and tyrosine have specific neutral losses of -H_3_O_4_P_1_. The use of labelling and reporter techniques in quantitative proteomics, such as TMT and iTRAQ, lead to even more extended sets of diagnostic ions and neutral losses required for comprehensive spectrum annotation^33^. Furthermore, there are classes of modifications with highly complex fragmentation like glycans and lipids. These latter categories of peptides are not yet widely supported in current database search and spectrum annotation tools^34,35^.

Apart from the complexities of fragmentation and post-translational modifications there is a multitude of other compounding factors in peptidoform annotation: ambiguous amino acids, unlocalized modifications, cross-linked peptides, and the possible occurrence of chimeric spectra. There are currently tools to annotate such individual cases^5–18^, but none that can handle all these complexities across the board and simultaneously within the same spectrum. Here we present a new tool, the Annotator, able to unify all these fields across peptides (bottom-up) and proteins (middle- and top-down). Based on the unified peptide/peptidoform notation ProForma 2.0^36^, Annotator allows users to quickly and interactively annotate mass spectrometry data from any of the listed fields. Additionally, the annotation code is available as a Rust and Python library allowing easy scripting for annotation statistics to support scoring of full proteomics-scale datasets.

## Results

### Rust library for ProForma-based peptidoform annotation

We aimed to develop an open-source graphical program for comprehensive spectrum annotation across a wide variety of proteomics applications. For this we developed the underlying open-source library rustyms to handle mass spectrometry peptide fragmentation data, written in Rust for speed and robustness. Rustyms uses ProForma 2.0 to define the peptides/peptidoforms.^36^ ProForma standardizes notation and definition of many complex peptide structures, including predefined modifications, glycans, and chemical cross-linkers, as illustrated in Figure 1. Rustyms can generate theoretical fragmentation data for any ProForma peptidoform and allows extensive user control over the fragmentation model.

Spectra can be loaded and annotated in a variety of ways. The user can import a raw spectrum file (supported file types are mgf, mzML, Bruker TDF, or Thermo RAW), navigate to the correct spectrum and provide the sequence manually. Additionally, many file types of identified peptides, for example mzTAB, are supported to correlate the peptide with the right spectrum, which allows for easier navigation through larger datasets. A universal spectrum index (USI)^37^ can also be used to access data from public repositories, which will download the right spectrum for annotation. The user can then specify the fragmentation model, which defines the theoretical fragmentation (see Figure 2A). This controls which backbone and satellite fragment series occur, where they occur, what neutral losses are expected, and which charges are allowed. Additionally, this model controls other fragmentation options, including glycan, diagnostic, and immonium ions. Models for multiple fragmentation modes are predefined, including CID/HCD, ETD, EThcD/ETcaD, EAD, EACID, and UVPD. In addition, custom models can be specified, saved and shared by the user. Additional settings for the annotation can then be defined, including mass tolerance and noise filter, and the ProForma definition provided. The annotation will result in a fragment overview on the peptidoform and the annotated spectrum.

**Figure 2.**
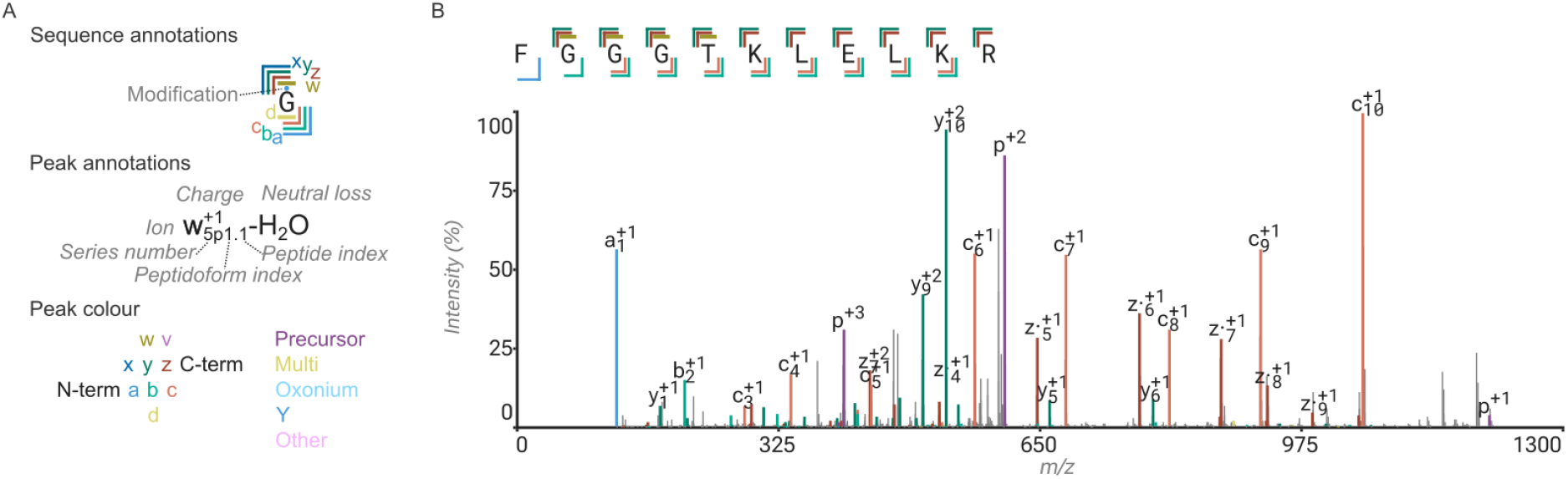
Peptide spectrum annotation with rustyms and ProForma in Annotator. A) Legend for display of fragments and modifications in Annotator. B) Example annotation of simple linear peptide of the light chain of mouse monoclonal antibody 139H2 by thermolysin digestion, analyzed by LC-MS/MS with EThcD fragmentation.

### Annotation of linear peptides and chimeric spectra

An illustrative example annotation of a peptide fragmentation spectrum by EThcD is shown in Figure 2. In this spectrum backbone bond cleavages are found with corresponding fragment ions at all positions for a sequence coverage of 11/11; 14% of all peaks in the spectrum are annotated, explaining 42% of all MS2 intensity, and matching 40% of all theoretically possible fragments. This in contrast to the annotation with a bare bones model, including only the expected b, c, y, z fragments, without any neutral losses, which results in a sequence coverage of 10/11 (due to the missing a1 ion), 11% of all peaks annotated, 38% of all MS2 intensity annotated, and 46% of all theoretical fragments found. In both cases, the vast majority of unannotated peaks correspond to the heavier isotopologs of the matched monoisotopic fragments. Additionally, Annotator provides a false match rate estimation for the fraction of annotated peaks and their intensity. In this example the false match rates are 4% of matched peaks, 6% based on intensity for the full model, or 3% (peaks) and 2% (intensity) based the bare bones model. This false match rate is determined by shifting the spectrum up and down in m/z by an arbitrary ± 25+π *Th* and taking the average fraction of peaks and their sum intensity, divided by the corresponding statistics for the untransformed spectrum.

Annotator additionally allows for the analysis of chimeric spectra, resulting from coisolation of peptidoform ion precursors in DDA data, and the common mode of operation in DIA or all-ion fragmentation techniques. Chimeric peptides can be expressed in ProForma using the two peptidoform sequences separated by a plus. An illustrative example of a chimeric peptidoform annotation can be seen in Supporting Figure S1. Annotator will display statistics for each separate peptidoform and for the combined total. These statistics can also be generated for more extensive datasets by using the ‘multi-annotator’ program distributed with rustyms, as further explained below.

### Annotation of PTMs and diagnostic ions

Annotator accommodates all PTMs deposited in PSI-MOD^38^, UNIMOD^39^, GNOme^40^, XLmod^41^, and RESID^42^, including diagnostic ions, and allows for additional custom definitions of modifications, neutral losses and diagnostic ions (which can be saved and shared between projects and users). Protein glycosylation is amongst the most frequent and abundant co/post-translational modifications, especially in human-derived samples (*i*.*e*. blood, tissue).^43,44^ Furthermore, these glycan modifications can be incredibly heterogenous, both in composition, as well as in structure, as many glycan isomers exist.^45,46^ In ProForma glycans can be defined in two major ways: as a composition of monosaccharides or as an identifier from the GNOme database.^47^

Depending on the fragmentation techniques used, glycopeptides give rise to characteristic diagnostic ions and neutral losses.^34,48^ With ETD (electron transfer dissociation) and low kinetic energy ECD (electron capture dissociation), glycans stay attached to the peptide and glycans can be localized on the peptide backbone by examining the mass shifts of the c- and z-ions. Using CID/HCD type fragmentation (including supplemental activation in electron-based fragmentation methods), the glycan moieties themselves can fragment too, leading to the formation of B-, Y-, and internal oxonium ions (see also Figure 1B). B-ions are the terminal fragments of the glycan, Y-ions are the fragments containing the entire peptide backbone with glycan core fragments attached, and internal oxonium ions are the fragments where multiple B/Y breakages have occurred. All these ions are of particular interest when trying to elucidate glycostructural elements from the MS2 data, for example to distinguish whether a fucose is localized on the core or on the branch.

Rustyms can generate B-, Y-, and internal oxonium ions for both glycan definitions. For structural glycans (*e*.*g*. using a GNOme identifier), all possible combinations of broken glycan bonds are identified, and the resulting list of glycan and glycopeptide fragments are annotated with their respective broken bonds. For glycans defined by composition (*i*.*e*. [Glycan:HexNAc3Hex4]), all unique combinations of monosaccharides are generated and used to annotate all possible B-, Y- and internal oxonium ions. The graphical annotation of glycopeptide spectra in Annotator includes the automated generation and use of Symbol Nomenclature for Glycans (SNFG) for B-, Y-, and oxonium ions. These unique features of Annotator may benefit the expanding glycoproteomics community, providing a versatile tool to investigate glycostructural features on glycopeptides, as further highlighted below.

Multiple possible glycan structures can be defined simultaneously to annotate within the same spectrum, such that the best supported structure can be identified from the available evidence, as illustrated in Figure 3A, where the position of a fucose moiety (branch vs core) is mapped onto an N-glycopeptide. The glycan composition (HexNAc(5)Hex(5)dHex(1)NeuAc(2)) identified by the Byonic database search is compatible with several possible structures listed in the GNOme database: G75079FY (depicted on peptide 1 in Figure 3A), G52512ZN (depicted on peptide 2 in figure 3A), and G19935MZ (not shown). These three structures differ in two major ways: 1) the fucose is localized either on the branch HexNAc or on the glycan core HexNAc, and 2) the 5^th^ HexNAc is either connected as a branch, or a bisecting GlcNAc. These structural differences may exhibit different biological effects, which makes it important to distinguish them. At first glance, it seems that both fucose positions are supported by the fragments, however, it is known that fucoses can “hop” in the gas phase on glycopeptide ions, from the core to the branch position.^49^ With the clearly annotated Y-ions (in blue), this spectrum provides strong support for a core-fucosylated structure. Furthermore, the fragment containing the bisecting GlcNAc is annotated (in blue), helping to assign this unique structural feature.

**Figure 3.**
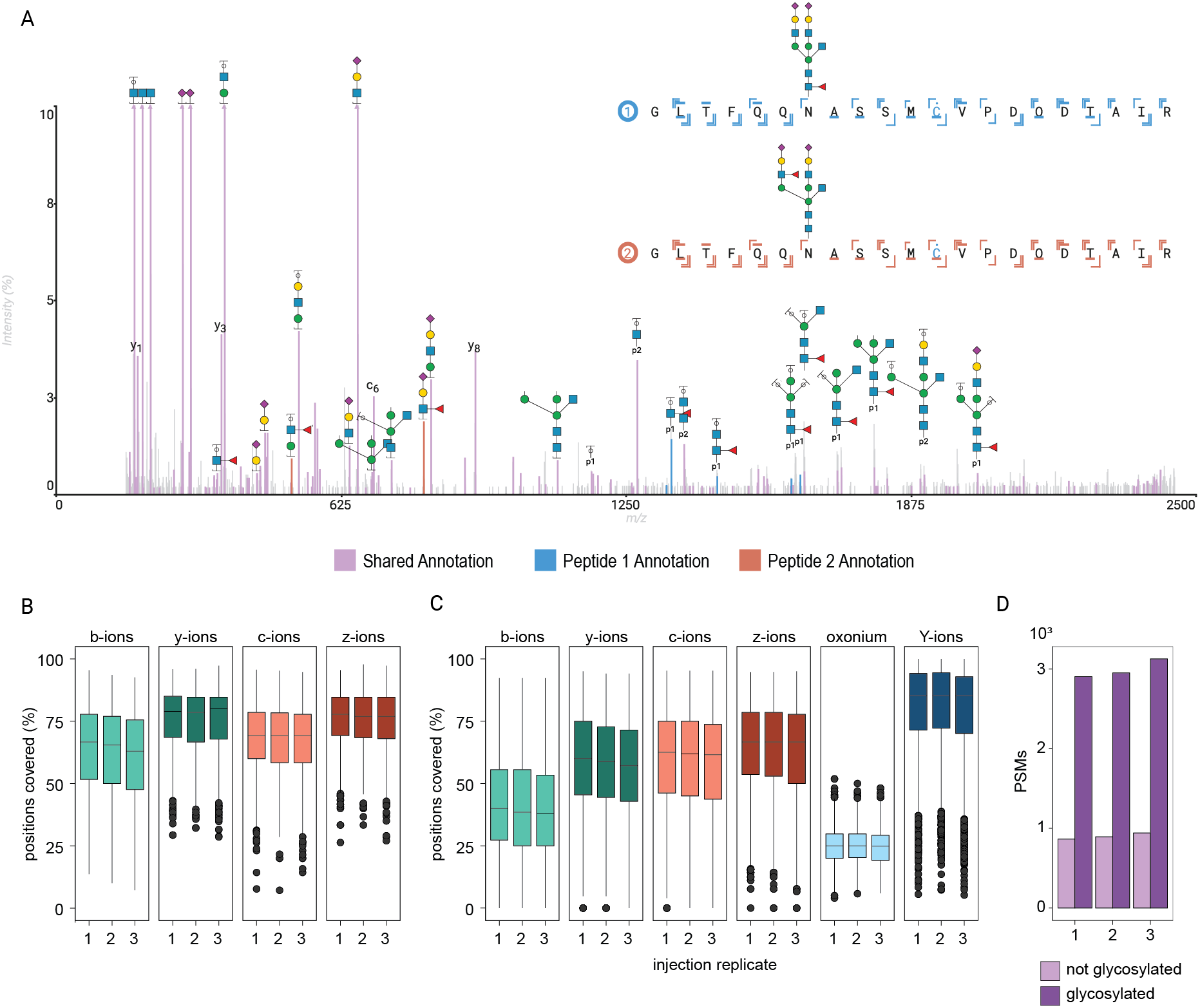
Annotation of post-translational modifications and diagnostic ions – structural analysis of glycopeptides. A) EACID spectrum acquired on the ZenoTOF 7600 of a tryptic IgM glycopeptide identified in human plasma with annotation of two compatible glycan structures to differentiate support for the two possible glycan isomers. The peaks in light purple color are the shared annotations of both structures, the blue and red peaks correspond to the unique fragments assigned to glycopeptide 1 and glycopeptide 2, respectively. B-D) The multi-annotator can extract ion sequence coverage in batch from the full plasma glycoproteomics dataset to evaluate fragmentation statistics for non-glycosylated PSMs (B) vs. glycosylated PSMs (C), with an overview of the total PSMs in the used dataset (D).

Apart from elucidating glycostructural elements, systematically going through ion coverages is beneficial for method optimization and quality control of different MS experiments. A utility distributed with rustyms, the multi-annotator, can parse through raw/mgf/mzML files and provide detailed ion coverages for each PSM. The generated output can be examined to adjust fragmentation parameters, or to assure consistent performance across runs and replicates, as shown in Figure 3B-D. Here, ion coverage of three injection replicates of complex samples (using ZenoTOF EACID data on glycopeptide enriched human plasma samples) are depicted, for non-glycosylated peptides (Figure 3B) and glycosylated peptides (Figure 3C), which demonstrated that the method used is robust and generates all the information required. Furthermore, it can be seen that y-ion coverage is lower for glycopeptides than for non-glycosylated peptides. For this analysis, each run had approximately 1000 non-glycosylated PSMs, and around 3000 glycosylated PSMs (Figure 3D).

Precise site localization is a crucial step of PTM analysis by mass spectrometry-based proteomics, which is also facilitated by Annotator. Protein glycosylation again serves as an illustrative example. While the localization of *N*-glycans is restricted to the *N*-glycan consensus sequence (N-X-S/T, where X is anything but proline), no such motif exists for *O*-glycans. *O-*glycosylation primarily occurs on threonine and serine residues, and many proteins have densely *O-*glycosylated regions, such as many viral proteins^50,51^, and mucins.^52^ One abundant human protein with such a domain is the immunoglobulin IgA1, which has multiple *O*-glycans within its hinge region, which all happen to co-localize on a single tryptic peptide.^53–56^ This large 38 amino acid peptide has up to 9 possible O-glycosylation sites. In Annotator, glycans can be moved around manually and sequence coverage can be assessed, until the highest number of backbone fragments are assigned, from which consequently the best location can be determined, as illustrated in Supporting Figure S2. This way Annotator can enable the formulation and use of delta scores for site localization like those used in phosphoproteomics^57–60^, by automating the procedure across a large dataset with the multi-annotator.

### Annotation of cross-linked peptides

Annotator also supports analysis of spectra beyond linear peptidoforms. Cross-linking MS has become an important tool in mass spectrometry-based structural biology and the analysis of protein interaction networks^61,62^. Due to the expanded search space for finding crosslinked pairs of peptides, the rates of false discovery might increase, making inspection and validation of crucial crosslinked peptide spectra an especially urgent and useful exercise in this application of proteomics. Cross-linked peptides provide another layer of complexity whose annotation is fully supported by the ProForma notation, including both chemical cross linkers and naturally occurring disulfide bonds. The annotation includes diagnostic ions and cleavable cross linkers. See Figure 4 for an example of the annotation of an HCD spectrum for a pair of peptides cross-linked to each other by using the cross-linker DSS.

**Figure 4.**
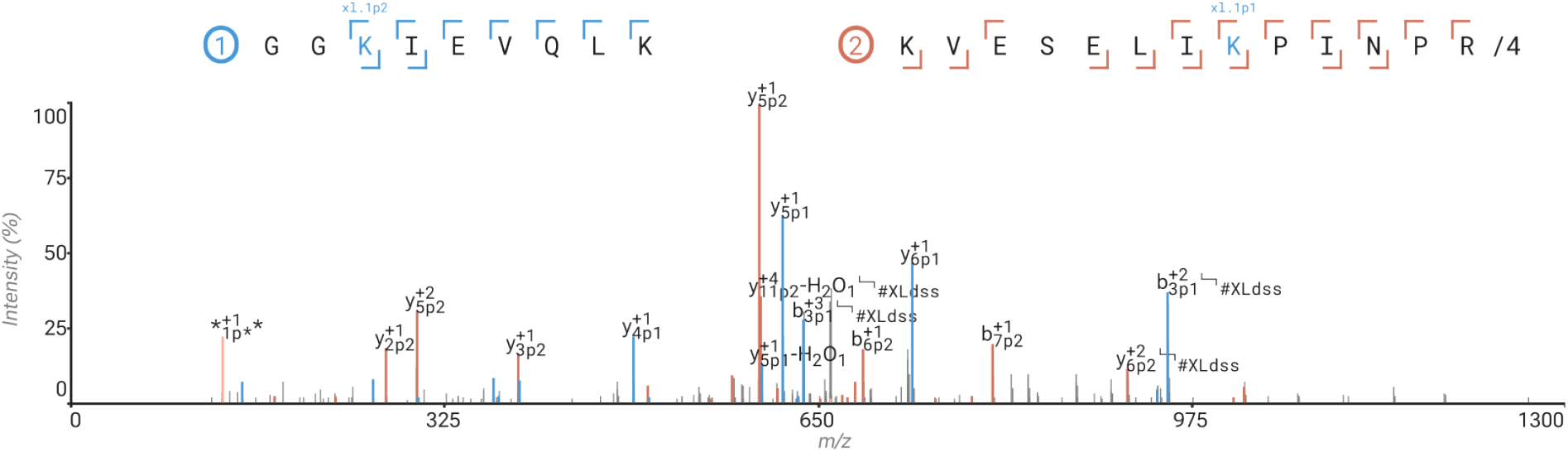
Annotation of cross-linked peptides. Cross-linked pair of tryptic peptides representing an intra-subunit link between the vitamin K-dependent protein S (ProS) subunit of the human C4b-binding protein complex (C4BP).

### Annotation of top- and middle-down proteomics spectra

Disulfide linked peptides represent an interesting case because they reflect the natural structure of folded proteins and often, multiple disulfide intra- and inter-links co-occur within and between multiple chains. This is especially relevant to middle- and top-down applications of proteomics, when intact proteins or sizeable proteolytic fragments thereof are analyzed. Such is the case for the antigen-binding fragment of IgG1 antibodies (Fab) where there are 5 disulfide bridges (see Figure 5A). The disulfide bridges are covered in ProForma and the Annotator with a custom modification, using formula H_-2_, with breakage: H_-1_:(empty), H_-1_:H_1_, H_-2_S_-1_:S_1_.^63^

**Figure 5.**
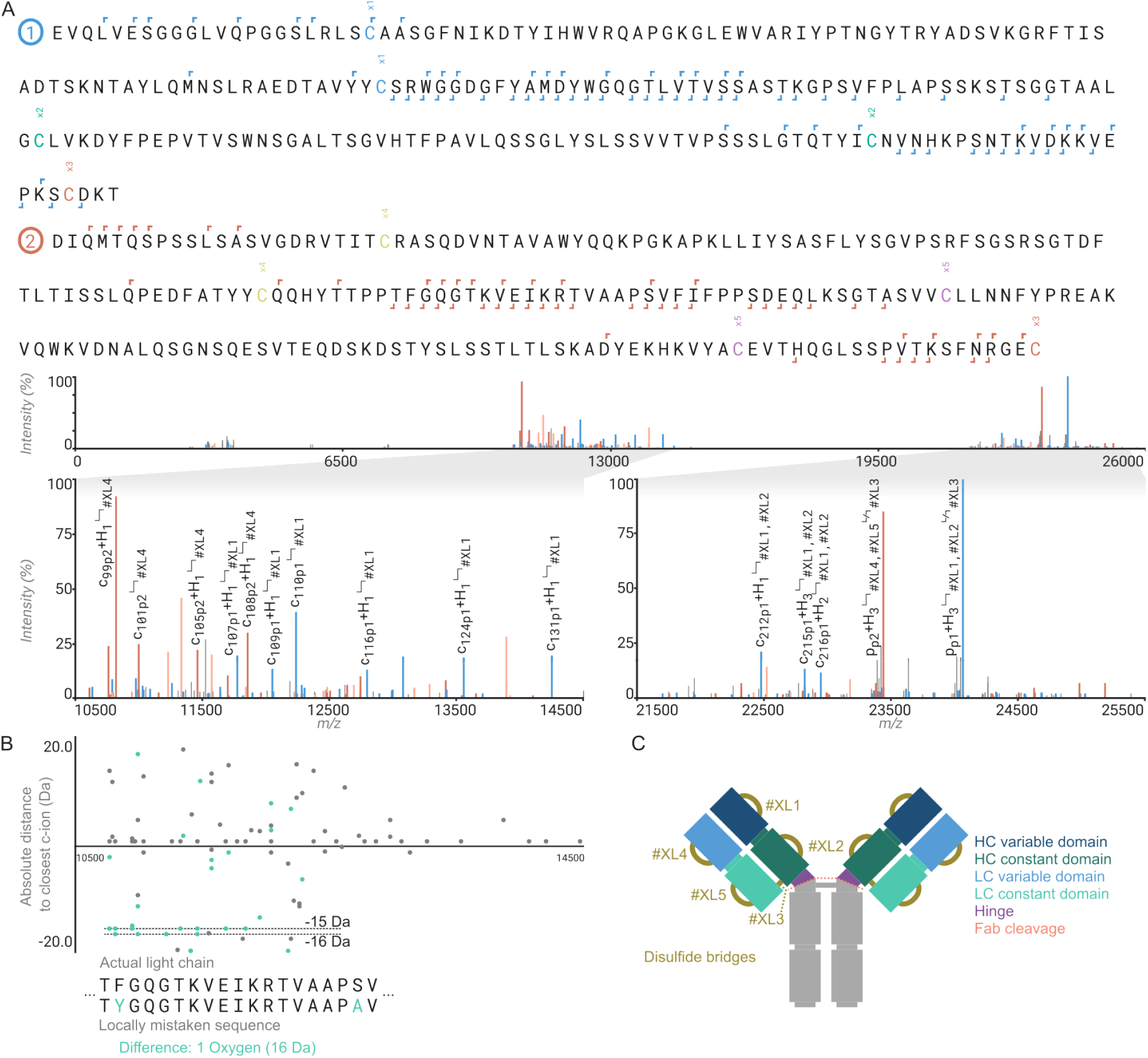
Annotation of disulfide linked protein chains for middle-down proteomics characterization of antibody fragments. A) Antigen binding fragment of the recombinant IgG1 Trastuzumab, showing variable domains of (1/blue) heavy chain and (2/red) light chain. Annotation is based on the deconvoluted top-down ETD fragmentation spectrum in the bottom panel. B) Error graph displaying the closest theoretical c-ion for all unannotated ions given the actual light chain sequence and a locally mistaken sequence. This highlights the use of this graph for detecting local errors in annotations. Also visible here is that big proteins can take up hydrogens as a result of ETD cleavage of disulfide bridges. C) A graphical representation of an antibody Fab, with the cross-links numbered as used in A.

Rustyms allows the combination of any of these features to fully be annotated within any complex MS2 spectrum (i.e., including disulfide cross linking and glycosylation or other PTMs within the same spectrum). Additionally, Annotator includes an error graph below the spectrum that can be used to inspect in detail a top-down spectrum to aid in *de novo* sequencing and validation of experimentally determined sequences of antibodies. This error graph displays the distance to the closest theoretical fragment for the selected ion series for all unidentified peaks. As depicted in Figure 5B this will show a range of peaks with a constant offset from the zero-line localized to the site of the misassigned residue in the sequence, allowing the manual refinement of a (top-down) *de novo* antibody sequence.

## Discussion

Rustyms and Annotator are comprehensive tools to analyze mass spectrometry-based proteomics data across a diverse range of applications. Several other peptide spectrum annotation tools are available, most of which will cover a specific subset of applications. Some of the preceding tools are focused on top/middle-down fragmentation, such as ExDViewer^5^, Omniscape^6^, and ProSight Light^7^. Annotator advances on these by allowing more complex peptide/peptidoform definitions as well as finer control over the annotation. Other tools have focused on fragmentation spectra in bottom-up proteomics, like FragmentLab^8^, LCMS Spectator^9^, Lorikeet^10^, Quetzal^64^, and mMass^11^. Annotator advances on these with more granular control over the fragmentation model and allowing more complex peptides/peptidoforms. Of these, mMass is notable for its support of cyclic peptides, which is not yet part of the ProForma and therefore not yet supported in rustyms and Annotator. Annotator also allows loading identified peptides from other software, including PEAKS^65^, Sage^66^, MaxQuant^67^, MSFragger^68^, Novor^69^, and O-Pair^70^, amongst others, in addition to Fasta and mzTAB^71^ files (as used by Casanovo^72,73^). Such features are also supported in FragmentLab, PeptideShaker^12^, and TOPPView^13^, although each of these has a different set of other software tools they support. Other libraries for handling mass spectrometry data have also previously been reported, including Open MS (c++)^14,74^, pyteomics (Python)^15,16^, and rforproteomics (R)^17^. Rustyms advances on these tools by allowing more complex peptide definitions. A project with similar goals is spectrum_utils (Python)^18,75^, which is a library allowing annotation and visualization with full support for ProForma 2.0; Annotator advances over this approach by having a graphical user interface.

We aimed to develop a universal peptide spectrum annotator. However, there are still some missing features that could be a good extension in the future. Firstly, cyclic peptides are not well supported in ProForma and as such are not well supported in Annotator. There are ways of defining them with cross-links, but this is not the best user experience and might not result in the expected fragmentation pattern. Secondly, internal fragmentation beyond immonium ions is currently not supported in Annotator, even though their use has attracted interest in top-down proteomics applications as of late^28–31^. Finally, the Annotator could be extended to create and load spectral libraries in mzSpecLib^76^ format to allow visualizing spectra from any annotation tool and to allow the annotations from the Annotator to be reused in other tools.

## Conclusions

Mass spectrometry-based proteomics covers a diverse range of applications and as such the complexity of the resulting data can be overwhelmingly large. With rustyms we present an open-source library for annotating a peptidoform of arbitrary complexity, and with the Annotator we present an open-source, easy-to-use graphical tool to visualize all these spectra. The combination of all layers of complexity in ProForma, ambiguous amino acids, modifications, modifications of unknown position, glycans, cross-linkers, and chimeric spectra allows Annotator to be used as a comprehensive tool across the wide range of mass spectrometry-based proteomics applications. Therefore, rustyms and Annotator will be in our view a welcome addition to the ever-expanding software toolbox for mass spectrometry-based proteomics.

## Methods

### Bottom-up glycoproteomics

N-glycoproteomics was performed on a pooled human plasma (VisuCon-F Normal Donor Set (EFNCP0125), Affinity Biologicals), digested with Trypsin and LysC, and subsequently enriched for glycopeptides using cotton-HILIC SPE^77^, as previously described ^78^.

For *O*-glycoproteomics, human colostrum IgA (Sigma Aldrich) was denatured, reduced and alkylated in SDC buffer (0.1 M TRIS/HCl pH 8, 40 mM TCEP, 100 mM CAA, 1% SDC, *v/v*), and subsequently digested with trypsin (1:100). Samples were desalted using OASIS HBL plates (Waters), according to manufacturer’s instructions. Peptides were dried and reconstituted in 0.1% TFA.

Data was acquired using an Ultimate 3000 HPLC (Thermo Fisher Scientific), coupled online to a ZenoTOF 7600 Mass Spectrometer (SCIEX). The glycopeptides were trapped on a PepMap™ Neo 5 μm C18 300 μm X 5 mm Trap Cartridge (Thermo Fisher Scientific). *N*-glycopeptides were separated on an Evosep Performance column (15 cm x 150 µm inner diameter, C18 stationary phase, 1.5 µm particle size, Evosep) which was kept at 55 °C, while the *O*-glycopeptides were separated on an Ion Opticks Aurora Elite XS column (15 cm x 75 µm inner diameter, C18 stationary phase, 1.7 µm particle size, IonOpticks) which was kept at room temperature. The LC mobile phases were water with 0.1% formic acid (solvent A) and 80% acetonitrile in water with 0.1% formic acid (solvent B) (both UPLC grade, Biosolve) and a constant flowrate of 3 μl/min was used.

For the *N-*glycoproteomics, the concentration of solvent B was kept constant for 1 min at 3%, after which it was gradually increased to 30% over 39 min, to 44% in 5 min, to 55% in 2 min and to 99% in 2 min, where it was kept constant for 5 min, after which it was decreased to 3% for 10 minutes. The ZenoTOF 7600 was equipped with the OptiFlow 1-50 μl source, and the orthogonal spray was used. The source settings were the following: curtain gas, 25; CAD gas, 7; Ion source gas 1, 12 psi; Ion source gas 2, 60 psi; source temperature, 150 °C; spray voltage, 3600 V. The MS1 parameters were: mass range, 700-2000 *m/z;* collision energy, 10 V; “Peptide” workflow; max candidate ions, 25; intensity threshold, 300 counts/s; charge states, 2-10, exclude former candidate ions for 9 s. For MS2: mass range, 150-3000 *m/z*, accumulation time, 0.045 s, dynamic collision energy activated (charge 2: slope 0.049, intercept –11, charge 3-10: slope 0.05, intercept –12); electron beam current, 7000 nA; electron KE, 9 eV, reaction time, 10 ms. While for the *O-*glycoproteomics, the concentration of solvent B was kept constant for 1 min at 3%, after which it was gradually increased to 40% over 38 min, to 55% in 5 min, and to 99% in 1 min, where it was kept constant for 5 min, after which it was decreased to 3% for 10 minutes. The ZenoTOF 7600 was equipped with the OptiFlow Nano, with source settings: curtain gas, 35; CAD gas, 7; nano cell temperature, 300 °C; nano gas 1, 10; spray voltage, 1500 V. The MS1 settings were: mass range, 400-2000 *m/z;* collision energy, 10 V; “Peptide” workflow; max candidate ions, 15; intensity threshold, 300 counts/s; charge states, 2-10, exclude former candidate ions for 4 s. For MS2: mass range, 150-3000 m/z, accumulation time, 0.065 s, collision energy, 12 V; electron beam current, 7000 nA; electron KE, 9 eV, reaction time, 20 ms.

Raw data files were searched using PMI-Byonic (v.5.5.2, Protein Metrics). The plasma samples were searched against a focused database of plasma proteins, the IgA against the tryptic peptide in IgA1 which contains all O-glycans (HYTNPSQDVTVPCPVPSTPPTPSPSTPPTPSPSCCHPRLSLHR). The N-glycan database consisted of 279 structures (max. 1 per peptide), the O-glycan database of 4 core 1 O-glycans (max. 7 per peptide). Carbamidomethylation of C was set as fixed modification, variable modifications included oxidation on M or W and pyroglutamic acid formation on protein and peptide N-terminal Q and E.

Subsequent analysis was performed in R, where data was filtered to 1% FDR. All glycan compositions were matched to possible GNO-structures. This was then used as input for multi-annotator to calculate sequence coverage of different ion types. Peak annotations were performed in the Annotator. Specific spectra shown here are picked randomly.

### Top-down spectra for the Fab of Trastuzumab

40 µL of CaptureSelect FcXL Affinity Matrix beads were added to a spin column and washed as per the manufacturer’s instructions. Trastuzumab (100 µg, kindly provided by Roche Penzeberg) in 150 µL phosphate buffer (PB) was incubated with the beads for 1 hour. The spin column was then washed twice with 150 µL PB and twice with 150 µL Milli-Q water (MQ). Digestion was performed overnight at 37°C with shaking (750 RPM) using 50 µL PB containing 10 µg of in-house produced r-IgdE. The column flow-through (FT) was collected after centrifugation.

The sample was denatured with 150 µL 8M guanidine at 60°C for 10 minutes (750 RPM), then buffer exchanged using 10 kDa cutoff Amicon centrifugal filters. Following manufacturer instructions, the filter was conditioned, and sequential rounds of solvent exchange were performed: 6 cycles of 400 µL each, including 1 round of 10% acetonitrile (ACN) with 1% formic acid (FA), 4 rounds of 20% ACN with 0.1% FA, and 1 final round of 30% ACN with 0.1% FA. Flow-through was discarded after each step. The final sample was eluted by inverting the spin column into a fresh tube and centrifuging at 1000 × g for 1 minute.

The sample was diluted to 1 µm and directly infused using a custom gold-coated glass capillary integrated into a Nanospray Flex™ Ion Source. The needle voltage was set to 880 V, with Sheath Gas, Auxiliary Gas, and Sweep Gas at 0 arbitrary units, and the source voltage at 15 V. The ion transfer tube temperature was maintained at 275 °C.

MS^2^ scans were performed, selecting the most abundant charge state for quadrupole isolation with a 3 m/z isolation width. The AGC target was 1000% with a maximum injection time of 100 ms. ETD activation was applied for 5 ms using a reagent target of 6e6. The Orbitrap operated at 120K resolution across a mass range of 1000–6000 m/z. The RF lens voltage was set to 60%, and each scan consisted of five micro scans.

44 MS^2^ scans were summed in Freestyle and deconvoluted with Xtract. The full m/z range was analyzed, outputting MH^+^with H^+^as the adduct. A charge range of 2–50 was specified, with a minimum of one detected charge.

## Supporting information

Example Spectra

## Data Availability

Both rustyms and the Annotator are open source on GitHub (https://github.com/rusteomics/mzcore and https://github.com/snijderlab/annotatorrespectively), dual licensed under MIT or Apache-2.0. The Annotator has prebuilt binaries for all major platforms on GitHub and is listed on the windows package registry, meaning it can be installed on any windows machine with the following command “winget install –id Snijderlab.Annotator”. The glycan and Fab fragmentation data was deposited to the ProteomeXchange Consortium via the MassiVE partner repository with the dataset identifier MSV000096837 with ProteomeXchange identifier PXD059727. The USIs (Universal Spectrum Identifiers) for all spectra can be found in Supporting Table S1.

## Supporting Information

- Supporting Data S1: Example MGF file with all spectra used in the figures
- Supporting Table S1: universal spectrum indices for all displayed spectra
- Supporting Figure S1: Annotation of multiple peptidoforms within chimeric scans
- Supporting Figure S2: Site-localization of post-translational modifications

## Acknowledgements

The authors would like to thank Andrea Trezza, Bastiaan de Graaf, Cynthia Kelley, Dina Schuster, Gabriela Koike, Laura Perez-Paneda, and Samiksha Sardana of the Biomolecular Mass Spectrometry group at Utrecht University for testing and guiding the design of the Annotator. This research was funded by the Dutch Research Council NWO Gravitation 2013 BOO, Institute for Chemical Immunology (ICI; 024.002.009) and the European Research Council Executive Agency HORIZON ERC-2022-STG (FLAVIR; 101077640) to J.S.

## Author contribution

D.S. developed the software. S.J. acquired the glycoproteomics data, R.L. acquired the top-down data on the Fab of trastuzumab. D.S took the lead in writing the manuscript. All authors helped steer the software and contributed to the final manuscript. A.J.R.H and J.S. provided supervision and funding for the project.

## For Table of Contents only

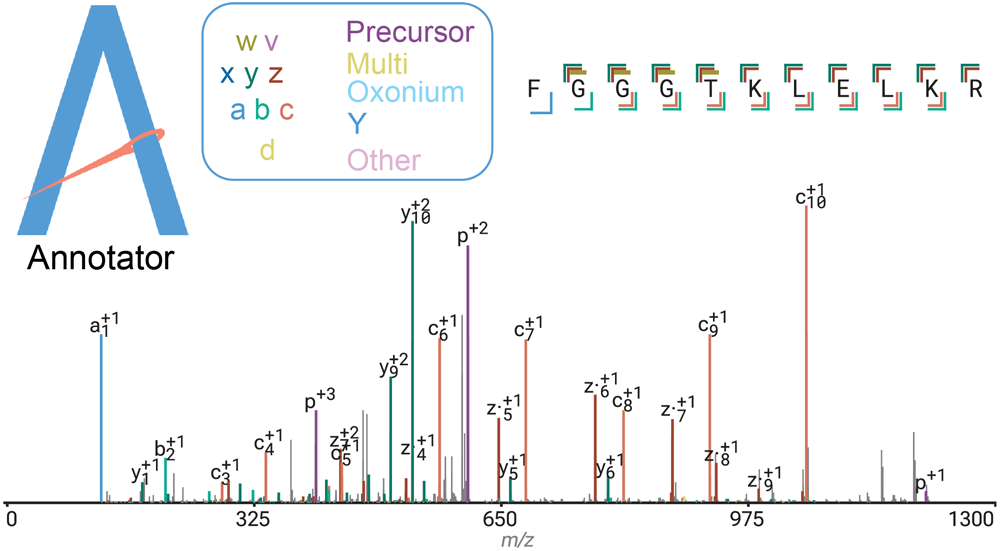

